# Characterization of a human-specific tandem repeat associated with bipolar disorder and schizophrenia

**DOI:** 10.1101/311795

**Authors:** Janet Song, Craig B. Lowe, David M. Kingsley

## Abstract

Bipolar disorder (BD) and schizophrenia (SCZ) are highly heritable diseases that affect over 3% of individuals worldwide. Genomewide association studies have strongly and repeatedly linked risk for both of these neuropsychiatric diseases to a 100 kb interval in the third intron of the human calcium channel gene *CACNA1C.* However, the causative mutation is not yet known. We have identified a novel human-specific tandem repeat in this region that is composed of 30 bp units, often repeated hundreds of times. This large tandem repeat is unstable using standard polymerase chain reaction and bacterial cloning techniques, which may have resulted in its incorrect size in the human reference genome. The large 30-mer repeat region is polymorphic in both size and sequence in human populations. Particular sequence variants of the 30-mer are associated with risk status at several flanking single nucleotide polymorphisms in the third intron of *CACNA1C* that have previously been linked to BD and SCZ. The tandem repeat arrays function as enhancers that increase reporter gene expression in a human neural progenitor cell line. Different human arrays vary in the magnitude of enhancer activity, and the 30-mer arrays associated with increased psychiatric disease risk status have decreased enhancer activity. Changes in the structure and sequence of these arrays likely contribute to changes in *CACNA1C* function during human evolution, and may modulate neuropsychiatric disease risk in modern human populations.

---

More than 3% of the global population has bipolar disorder (BD) or schizophrenia (SCZ), and both diseases are among the top 25 causes of disability worldwide (Saha et al., 2005; Merikangas et al., 2011; GBD 2015 Disease and Injury Incidence and Prevalence Collaborators, 2016). Along with the disability cost, both disorders are associated with an increased risk of suicide (Krishnan, 2005; Baldessarini et al., 2006). There are limited treatment options for BD and SCZ, and the burden of these diseases may be increasing (Saha et al., 2007). Improved diagnosis and treatments may come from a better understanding of the molecular pathways that contribute to disease risk.

Both BD and SCZ are highly heritable. While they are classified as different diseases based on their clinical symptoms, they share a similar set of genomic risk variants (Forstner et al., 2017). Genome-wide association studies (GWAS) for BD and SCZ have consistently implicated risk variants in or near genes involved in calcium signaling (Ferreira et al., 2008; Ripke et al., 2011; Sklar et al., 2011; Smoller et al., 2013; Ripke et al., 2013, 2014; Ruderfer et al., 2014). Calcium signaling-related genes are also enriched for rare variants in families multiply affected by BD (Ament et al., 2015) and in individuals with SCZ (Purcell et al., 2014; Andrade et al., 2016), suggesting that calcium signaling plays an important role in both BD and SCZ etiology.

Some of the strongest and best-replicated associations for BD and SCZ map within the *CACNA1C* gene, which encodes the pore-forming subunit of the Ca_V_1.2 calcium channel (Dedic et al., 2018). Disease-associated single nucleotide polymorphisms (SNPs) are in strong linkage disequilibrium with each other and contained within a 100 kb region of the gene’s third intron (Ferreira et al., 2008; Ripke et al., 2011; Sklar et al., 2011; Smoller et al., 2013; Ripke et al., 2013, 2014; Ruderfer et al., 2014; Nie et al., 2015). Underscoring the importance of this genomic region for psychiatric disease in humans, genotyped SNPs at this locus have also been associated with anxiety, depression-related symptoms, obsessive-compulsive symptoms, decreased performance in memory-related tasks, major depression, and autism (Erk et al., 2010; Bigos et al., 2010; Green et al., 2010; Casamassima et al., 2010; Liu et al., 2011b; Hori et al., 2012; Zhang et al., 2012; Smoller et al., 2013; He et al., 2014; Li et al., 2015).

The causative variants at loci identified by GWAS could be the assayed SNPs themselves (Guenther et al., 2014), or other variants tightly linked to the SNP markers (Claussnitzer et al., 2015). Previous studies have investigated the functional consequences of the genotyped SNPs in *CACNA1C* and other closely linked SNPs (Roussos et al., 2014; Eckart et al., 2016). However, the mutation responsible for the association between SNPs within *CACNA1C* and human neuropsychiatric diseases is still unknown.

Given the difficulty in identifying causal mutations at *CACNA1C* over the past ten years (Ferreira et al., 2008), we considered whether there might be additional structural variants at the locus that are not easily detected using current genotyping and sequencing methods. For example, copy number variants and expansions and contractions of micro- and mini-satellite sequences can be difficult to identify with short-read or Sanger sequencing technologies. Nevertheless, these types of mutations have been implicated in a wide range of neurological diseases, including Huntington’s disease, spinocerebellar ataxia, and fragile X syndrome (Verkerk et al., 1991; The Huntington’s Disease Collaborative Research Group, 1993; Lindblad et al., 1996; Renton et al., 2011; DeJesus-Hernandez et al., 2011).

To search for unrecognized copy number variants at the *CACNA1C* locus, we examined regions of the genome where no mutations were identified by large-scale sequencing projects such as the 1000 Genomes Project (1000 Genomes Project Consortium, 2015), yet DNA sequencing reads consistently differed from the reference human assembly. We identified one such region (hg38; chr12:2255791-2256090) within the 100 kb interval associated with BD and SCZ. In the most recent human reference genome (hg38), this region is assembled as a tandem repeat composed of ten 30 bp units. Chimpanzees and other non-human primates have a single instance of a homologous 30 bp sequence at this location (Fig. 1A), suggesting that an ancestral 30-mer sequence has expanded in the *CACNA1C* intron during human evolution. Strikingly, the number of reads from individuals in the 1000 Genomes Project that map to this 300 bp segment is 3-379x greater than expected based on the reference assembly, and these reads contain multiple base substitutions. Further investigation identified a longer (3.3 kb) repeat array at this location in the Venter assembly (HuRef) (Levy et al., 2007) and an approximately 6 kb repeat array in the genome of a hydatidiform mole sequenced to 40x coverage with long-read technology (Chaisson et al., 2015). Collectively, these data suggest that humans have a large and variable tandem repeat in the neuropsychiatric risk-associated region of *CACNA1C.* The size of the tandem array is likely under-represented in the human reference genome by one or two orders of magnitude based on empirical estimates from read depth coverage (see Supplemental Methods).

**Figure 1:**
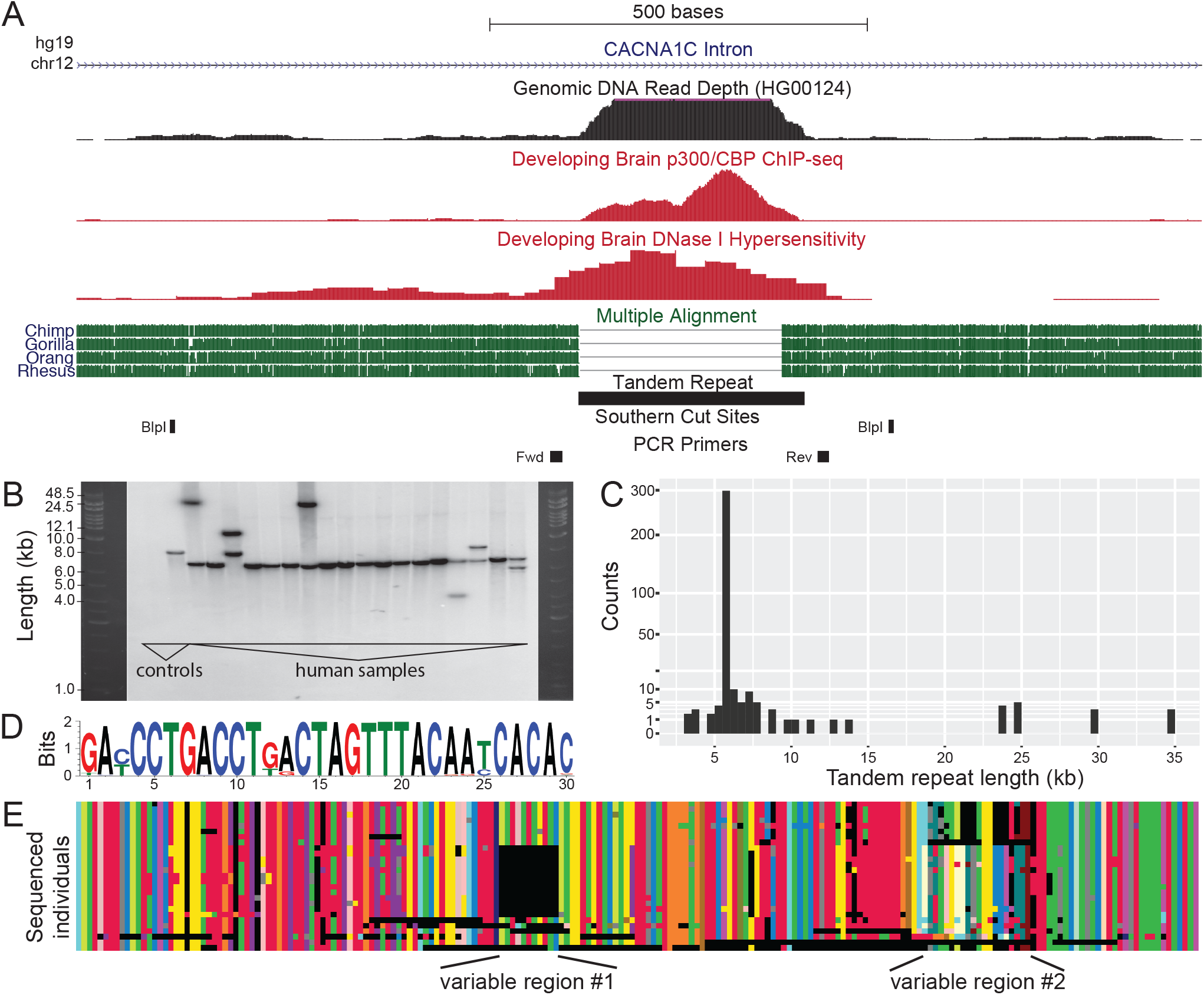
Human-specific tandem repeat region is composed of 30-mer sequence units repeated head-to-tail in multi-kilobase arrays. (A) The tandem repeat is located in the third intron of *CACNA1C.* Chimpanzee and other apes have a single copy of the 30 bp segment, while the human reference assembly predicts 10 copies of the 30 bp segment. There is an abnormally large number of genomic DNA sequencing reads mapping to the tandem repeat region, consistent with this repeat being further expanded in human individuals. The repeat region also shows enrichment for p300/CBP binding and DNase I hypersensitivity in the developing human brain. (B) We performed Southern blot analysis on 18 human individuals by probing for the 30 bp repeat after digesting with BlpI. We also included two controls: mouse DNA (no orthologous sequence) and the 8kb vector from which the probe was transcribed. The human reference genome predicts a BlpI fragment of approximately 900bp. In contrast, all humans tested show much larger BlpI fragment sizes (4,000 to 35,000 bp), and many individuals show dual bands indicating distinct alleles at the locus. (C) Frequency distribution of 362 repeat allele lengths detected by Southern blot analysis. (D) The 30-mer sequence logo calculated from the 30-mer variants present in human repeat arrays that were sequenced with long-read (PacBio) technology. Some positions are nearly invariant, whereas others vary from 30-mer to 30-mer. (E) Structure and composition of tandem repeat arrays sequenced by PacBio long-read technology. Each row represents a different sequenced array, and each color represents a distinct 30-mer variant. Black regions indicate gaps that we have introduced to maximize repeat alignments between arrays. Many regions are organized similarly in all arrays, but common variable regions distinguish array subtypes.

To further characterize the size of the tandem arrays using independent methods, we examined DNA from humans and our closest living relative, the chimpanzee. Polymerase chain reaction (PCR) amplification and sequencing from six chimpanzees confirmed a single instance of a 30-mer sequence, which exactly matches the chimpanzee reference genome. In contrast, when we performed Southern blots on human DNA (see Supplemental Methods), we found restriction fragment sizes consistent with repeat arrays of 3,000 to 30,000+ bp, with the majority of human repeat arrays showing sizes of approximately 6,000 bp (Fig. 1B,C). We never observed a band size consistent with the human reference genome (hg38). The smallest band size seen in the 181 human samples we assayed (362 alleles) was 10 times larger than the repeat size annotated in the reference assembly (300 bp), while the largest was more than 100 times larger.

To understand why the human genome assembly appears to have a version of the tandem repeat that is not representative of the human population, we examined four bacterial artificial chromosome (BAC) clones derived from a single individual that were used in the sequencing and assembly of the human genome (Osoegawa et al., 2001). One BAC clone matched the length and sequence present in the assembly (300 bp). A BAC library made from a single individual should have at most two alleles; however, the four BACs all gave different tandem repeat lengths. Compounding this anomalous result, colonies picked from a single BAC clone are expected to be identical, but two of the four BACs produced subclones with varying tandem repeat lengths (Fig. S1). In our experience, multi-kb tandem repeats, whose size was determined by Southern blot, reduced in length after amplification by PCR under routine conditions, or when propagated using standard circular vectors in bacteria. We propose that the human reference assembly is based on a BAC clone that was correctly sequenced and assembled, but that the sequence present in the BAC is an artifact of the instability of this tandem repeat when cloned and propagated using standard methods. Given the large size disparity between the version represented in the current human genome assembly and the alleles detected by Southern blot, as well as the instability of this region, we believe that the allele present in the current human genome assembly is not present in humans.

In addition to variation in the length of this repeat region in the human population, the 30-mer units that comprise each array also show sequence changes. For example, the array in the reference assembly is composed of four identical 30-mer units and six unique units that each contain a small number of SNPs. This variability in 30-mers is also seen in the large number of reads from the 1000 Genomes Project that map to this area. To better understand this variation in tandem repeat arrays, even for arrays of the same length, we performed long-read (PacBio) sequencing of repeat arrays amplified from 20 individuals using optimized PCR conditions (see Supplemental Methods). The size of the resulting PCR fragments using our optimized conditions matched the corresponding repeat lengths determined from Southern blots. The sequenced arrays were entirely composed of 30-mer units repeated head-to-tail. Some positions in the 30-mer unit appear to be largely invariant (e.g. position 2 is almost always an A), whereas other positions are more variable (Fig. 1D, Fig. S2). For instance, the most common 30-mer unit (31%) is GACCCTGACCTGACTAGTTTACAATCACAC and the second most common (17%) is GATCCTGACCTGACTAGTTTACAATCACAC (difference underlined). When aligning tandem repeat array variants, the structural organization of the 30-mer units within each array emerged (Fig. 1E). Across all PacBio-sequenced repeat arrays, certain regions, such as the beginning and the end of each repeat array, contain the same 30-mer units organized almost identically. However, other regions (marked in Fig. 1E) are more variable and contain specific patterns of 30-mer units that are consistently found in only a subset of the sequenced arrays.

The presence of a large and variable repeat region in the third intron of the human *CACNA1C* gene raises the possibility that variation in the tandem array contributes to functional changes at the locus. To test whether the length or sequence of the tandem repeat region shows any association with genomic risk markers for BD and SCZ, we examined whole genome sequence reads from individuals in the 1000 Genomes Project. We limited our analysis to individuals of European or East Asian descent, the two groups in which BD and SCZ risk status has been previously associated with four SNPs clustered in the third intron of the gene (rs2007044, rs1006737, rs4765905, and rs4765913; Fig. 2A) (Ferreira et al., 2008; Ripke et al., 2011; Sklar et al., 2011; Smoller et al., 2013; Ripke et al., 2013, 2014; Ruderfer et al., 2014; Nie et al., 2015). We first identified all sequencing reads from this repeat region. To infer the length of the repeat array (average of the individual’s two alleles), we used the fraction of all sequencing reads for the individual that are from the repeat region (see Supplemental Methods). To estimate the sequence composition of the repeat array (averaged over the two alleles), we calculated the fraction of all 30-mers in the sequence reads identical to each observed sequence variant of the 30-mer unit (see Supplemental Methods). Since our length and composition statistics represent a mixture of the two alleles present in each person, we limited our analysis to individuals that are homozygous risk or protective at each of the four SNPs commonly associated with BD and SCZ (Fig. 2A). These SNPs are all tightly linked and define risk and protective haplotypes, making it possible to study repeat structures associated with risk or protective genotypes at *CACNA1C.*

**Figure 2:**
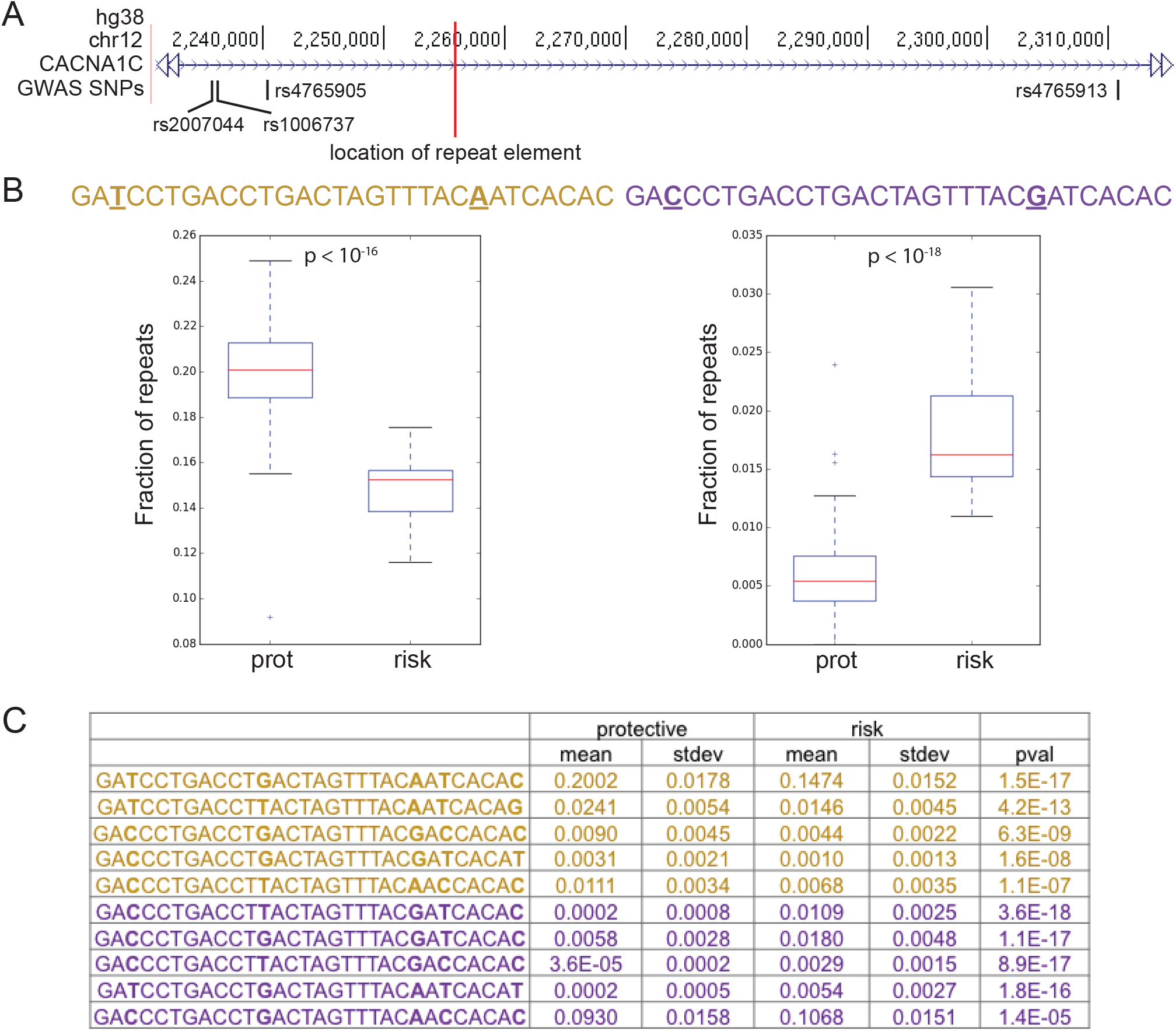
30-mer repeat variants are associated with protective or risk status at GWAS SNPs linked to neuropsychiatric disease. (A) Genome browser view of the third intron of *CACNA1C.* A red line marks the location of the repeat region. The human-specific 30-mer repeats are embedded in a region defined by four SNPs that are repeatedly associated with BD and SCZ. (B) We identified individuals from the 1000 Genomes Project that have the protective genotype at all four GWAS SNPs (protective haplotype) and individuals with the risk genotype at all four GWAS SNPs (risk haplotype). We only used European and East Asian individuals because GWAS have only been done with these populations. For each possible 30-mer repeat unit, we determined what fraction of 30-mers in the reads that map to this locus in each individual exactly match that particular variant. The 30-mer sequence on the left is significantly associated with the protective haplotype (prot), whereas the 30-mer variant on the right is significantly associated with the risk haplotype (risk). Base pair differences between the two 30-mer variants presented here are underlined. (C) The table lists the mean and standard deviation of the fraction of reads that exactly match a given 30-mer for individuals with the protective or risk haplotype. Repeats enriched in the protective haplotype group are shown in yellow, and repeats enriched in the risk haplotype group are shown in purple. The p-values were calculated using the Wilcoxon rank-sum test with Bonferroni correction (see Supplemental Methods).

We first tested whether repeat length is consistently associated with genotype status at the four GWAS SNPs. None of the four SNPs show a significant association with repeat length and the direction of effect is not consistent (Fig. S3). It does not appear that repeat array length is associated with the protective or risk genotypes at the GWAS SNPs, at least not in a simple manner.

We then tested whether specific sequence variants of the 30-mer unit are associated with the risk or protective alleles at the four GWAS SNPs. For each sequence variant of the 30-mer unit, we tested if its propensity to appear in reads from this repeat region differs between individuals that are homozygous for risk or protective genotypes at the four SNPs (Fig. 2B). We identified a number of 30-mer units that are consistently associated with a genotypic class across all four SNPs (Fig. S4). When considering only individuals that are homozygous protective at all four GWAS SNPs (protective haplotype) or homozygous risk at all four GWAS SNPs (risk haplotype), five 30-mer variants are associated with the protective haplotype, and five 30-mer variants are associated with the risk haplotype (Fig. 2C). These particular 30-mer units tend to be located in the variable regions observed when aligning the PacBio-sequenced repeat arrays (marked in Fig. 1E). Thus, while there is no straightforward association between the overall length of repeat arrays and the risk or protective haplotype, the abundance of particular 30-mer units is significantly and consistently associated with markers for psychiatric disease risk at *CACNA1C.*

The repeat region gives a significant signal for p300 enrichment in ChIP-seq experiments performed on tissue from the developing human brain (Visel et al., 2013), and also shows an open chromatin signal during human brain development (Kundaje et al., 2015) (Fig. 1A, Fig. S5). Both of these results are consistent with the repeat region acting as a distal enhancer element during brain development. To experimentally test whether the 30-mer repeat arrays show enhancer activity in developing neural cells, we cloned the single 30-mer sequence found in chimpanzees (30 bp), as well as 21 different human repeat arrays (3.5-6 kb), upstream of a basal promoter and a luciferase reporter gene (Fig. 3A). We used a linear cloning vector that greatly improved repeat stability, and we confirmed clone stability via comparison to the expected size and sequence as determined from Southern blot analysis and PacBio sequencing, respectively (see Supplemental Methods). We then transfected each construct into a human neural progenitor cell line (ReNcell Cx) and measured luciferase activity (see Supplemental Methods). The chimpanzee construct, containing a single 30 bp unit, weakly enhanced luciferase activity relative to the empty vector (*p* = 0.01), while the much larger repeat arrays found in humans significantly enhanced luciferase activity compared to both the empty vector and the chimpanzee construct (p < 10^−8^, Fig. 3B). These results suggest that genomic changes at the *CACNA1C* locus during human evolution have strengthened an existing enhancer element.

**Figure 3:**
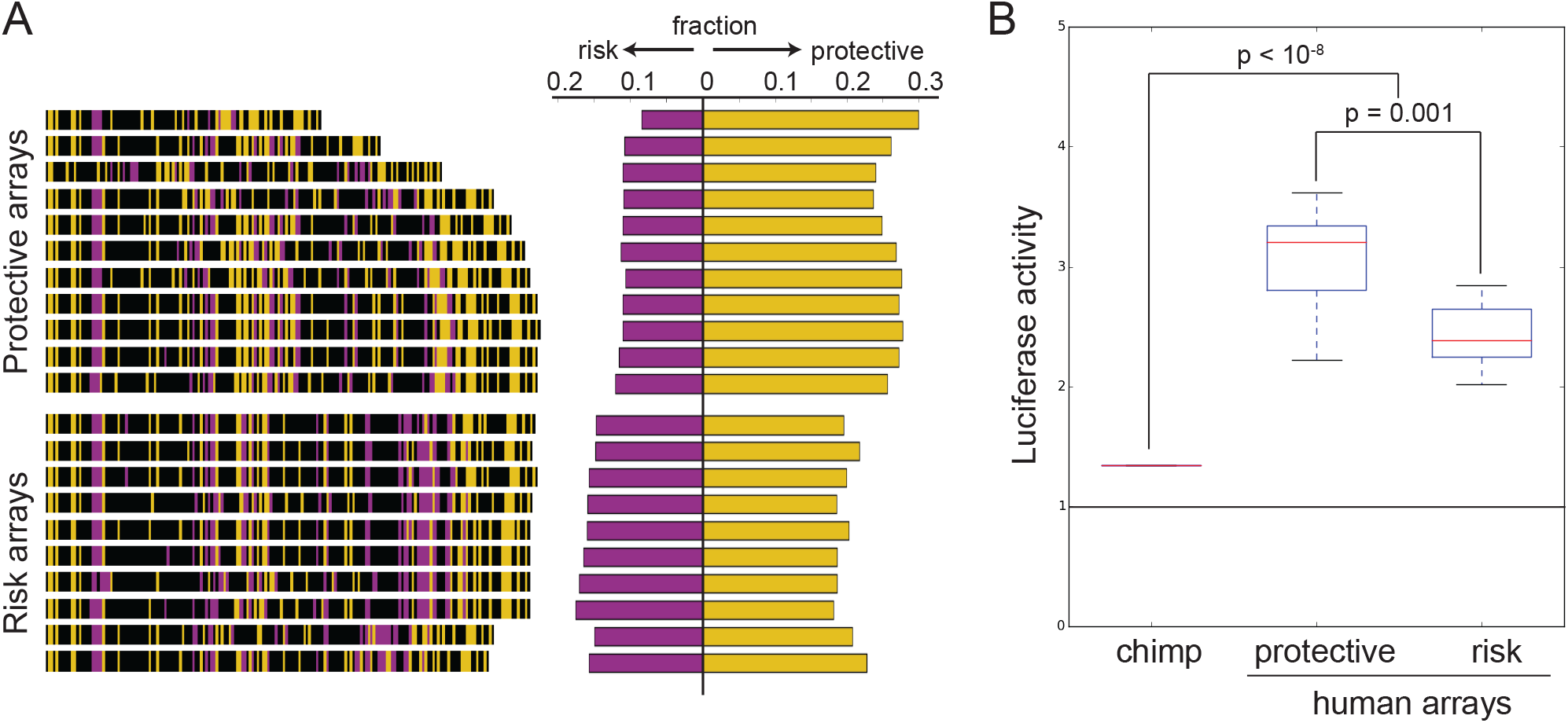
Human-specific repeat arrays act as enhancers in neural cells. The single 30-mer found in chimpanzees (30 bp) and twenty-one different human repeat arrays (3.5-6kb) were cloned upstream of a minimal promoter driving expression of the luciferase reporter gene. 30-mer variants significantly associated with the protective haplotype are colored yellow, 30-mer variants significantly associated with the risk haplotype are purple, and non-significant variants are black. The fraction of total 30-mer variants associated with either the risk or the protective haplotype varies between the protective-associated and risk-associated repeat arrays as expected, although the differences are subtle. (B) Constructs were assayed for luciferase activity in a human neural progenitor cell line (ReNcell Cx), as described in Supplemental Methods. Human repeat alleles drove significantly higher luciferase activity compared to the single 30-mer found in chimpanzees (*p* < 10^−8^). Protective arrays drove significantly higher luciferase activity than risk arrays (*p* = 0.001). The p-values were calculated using the Wilcoxon rank-sum test.

Although all of the tested human repeat arrays consistently acted as enhancers, there was substantial quantitative variation in enhancer strength among human repeat arrays (Fig. S6). To test whether enhancer strength varied for arrays linked to protective or risk GWAS SNPs for neuropsychiatric disease, we determined the genotypes of the individuals from which these human repeat arrays were cloned. Repeat arrays derived from individuals with the protective haplotype were classified as protective, and repeat arrays derived from individuals with the risk haplotype were classified as risk. For repeat arrays derived from individuals who are heterozygous at the GWAS SNPs, we determined the proportion of protective- and risk-associated 30-mer variants from PacBio sequencing. We then asked whether these proportions most closely resembled individuals with the protective haplotype or individuals with the risk haplotype in the 1000 Genomes Project and designated the ambiguous repeat arrays accordingly (Fig. S7, see Supplemental Methods). The repeat arrays characteristic of the protective haplotype drove significantly higher luciferase activity than repeat arrays characteristic of the risk haplotype (*p* = 0.001, Fig. 3). These data show that compositional differences between human repeat arrays lead to functional differences in enhancer activity, and suggest that differences in the repeat arrays may be causative genomic changes underlying the association between linked *CACNA1C* markers and susceptibility to neuropsychiatric disease.

Previous studies have tested whether risk and protective genotypes at the *CACNA1C* locus lead to higher or lower *CACNA1C* gene expression in the brain. Studies in the dorsolateral prefrontal cortex and cerebellum reported decreased *CACNA1C* expression in individuals with risk variants at human GWAS SNPs (Bigos et al., 2010; Gershon et al., 2014). In contrast, studies in the superior temporal gyrus and fibroblast-derived induced human neurons reported increased *CACNA1C* expression in individuals with risk variants at human GWAS SNPs (Yoshimizu et al., 2015; Eckart et al., 2016). Our data show that risk-associated repeat arrays have reduced enhancer activity in the particular human neural progenitor cell line we tested. We note that differences in human repeat arrays could also underlie more complex expression differences at other tissues or developmental timepoints. The base pair changes seen in particular 30-mer motifs that are associated with risk or protective genotypes alter the predicted binding sites for a number of potential trans-regulatory factors (Table S1). These factors themselves vary in expression and abundance in different brain regions (Bulfone et al., 1999; Lein et al., 2007; Liu et al., 2011a; Hawrylycz et al., 2012; Miller et al., 2014), which could in turn lead to differential effects of repeat variants at different times or places *in vivo.*

Previous studies of coding region mutations suggest that both loss-of-function and gain-of-function alterations in *CACNA1C* can lead to behavioral changes in mice and humans with similarities to BD and SCZ. For example, in mouse models where *CACNA1C* expression levels are either globally reduced or ablated only in specific brain regions, mice display increased anxiety and depression in behavioral tests such as the elevated plus maze, light-dark box, and learned helplessness test (Jeon et al., 2010; Dao et al., 2010; Lee et al., 2012; Dedic et al., 2018). Additionally, gain-of-function mutations in *CACNA1C* lead to Timothy Syndrome (TS) in humans, an autosomal dominant disease where afflicted individuals display autism-like symptoms in addition to a host of non-neurological patholo gies (Splawski et al., 2004, 2005). Although TS is normally lethal in young children, a rare TS patient who survived into his late teens developed BD (Gershon et al., 2014). These studies suggest that modulating *CACNA1C* expression levels, such as through human variation at the repeat arrays we report in this study, could result in behavioral changes associated with BD and SCZ.

We note that the 30-mer repeat arrays might have additional functional effects beyond the enhancer activities we characterize here. For example, the most common 30-mer sequences have open reading frames in both directions (Fig. S8A). Previous studies have shown that some tandem repeats are transcribed and translated even in the absence of conventional ATG start codons (Zu et al., 2011; Cleary and Ranum, 2013; Banez-Coronel et al., 2015). The tandem repeat also contains canonical splice site consensus sequences, including a donor site, an acceptor site, branch sites, and a polypyrimidine tract (Fig. S8B). Intriguingly, the single 30-mer found in chimpanzees has an A at the 17th position, whereas the vast majority of human 30-mers (99.94%) have a G at that position. This single base pair difference means that chimpanzees do not have canonical splice donor or acceptor sites at this locus. Finally, when organized in head-to-tail fashion, the 30-mers also form a CpG site located between the C that ends most 30-mers and the G that begins the next 30-mer (Fig. S8B). The tandem repeat arrays may affect translation, splicing, or methylation, in addition to forming a functional enhancer sequence within *CACNA1C*.

Our studies have identified a dramatic expansion of a 30-mer sequence that generates human-specific tandem arrays in a key gene related to calcium signaling, gene expression, and behavior. The human-specific repeat arrays show enhancer activity in human neural progenitor cells, and risk-associated versions of the tandem repeat have less enhancer activity than protective-associated versions. We hypothesize that generation of these repeat arrays has modified Ca_V_1.2 function during human evolution, and that structural and compositional differences of the 30-mer repeats among humans represent causal genomic changes that modify risk of neuropsychiatric disease in modern populations.

Many diseases that are particularly common in human populations occur at body locations that have also undergone dramatic and relatively recent evolution in the human lineage. For example, humans have a high incidence of lower back, knee, and foot problems, likely due to the recent evolutionary transition to upright bipedal walking (Pennisi, 2012). Over 70% of young adults develop impacted third molars (wisdom teeth), likely due to evolutionary reduction of jaw size in the human lineage (Stedman et al., 2004; Mann, 2013). Similarly, the high prevalence of neurological diseases in modern humans may be, in part, due to recent evolutionary changes in genes controlling brain size, connectivity, and function in humans compared to other primates (Oksenberg et al., 2013). The generation of new cellular and animal models that carry either chimpanzee or human 30-mer repeat arrays should make it possible to further characterize both the evolutionary and disease effects of this repeat region. In addition, the sequence differences in the 30-mer repeats can now be used as a novel feature to group patients into different genetic subtypes. Further stratification of patients based on *CACNA1C* repeat genotypes may prove useful for refined disease association studies, or for identifying cohorts that show differential response to drugs targeting calcium channel activity. Finally, our research illustrates how characterizing hidden variation in the human genome can uncover novel variants associated with both human evolution and disease.

## Description of Supplemental Data

Supplemental Data contains the Supplemental Methods, eight figures, and one table.

## Acknowledgements

We wish to thank members of the Kingsley Lab for useful discussions and comments on the manuscript. Research reported in this publication was supported in part by the NIDCR of the National Institutes of Health (K25DE025316, CBL), and by a National Science Foundation Graduate Research Fellowship and a Stanford Graduate Fellowship (JS). DMK is an Investigator of the Howard Hughes Medical Institute. DNA or tissue samples were obtained through the NIH Neurobiobank from the Human Brain and Spinal Fluid Resource Center (VA West Los Angeles Healthcare Center), the University of Maryland Brain and Tissue Bank, the Harvard Brain Tissue Resource Center, the University of Miami Brain Endowment Bank, the Mt. Sinai Brain Bank, and the Brain Tissue Donation Program at the University of Pittsburgh. The content is solely the responsibility of the authors and does not necessarily represent the official views of the National Institutes of Health.

